# Toward a digital twin for *Enterococcus faecium*: An experimentally validated manually curated genome-scale metabolic model of strain TX0016 reproduces high-throughput phenotyping data

**DOI:** 10.64898/2026.05.01.720924

**Authors:** Dina Samir Rasmi, Jayanth Krishnan, Yomna A. Hashem, Bernhard O. Palsson, Mona T. Kashef, Jonathan M. Monk, Ramy K. Aziz

## Abstract

Enterococci are Gram-positive opportunistic pathogens responsible for a wide range of nosocomial infections. One enterococcocal species, *Enterococcus faecium*, is steadily increasing in prevalence and has been listed among major multidrug-resistant ESKAPE pathogens. To gain systems-level insights into its metabolism and support discovery of potential therapeutic targets, we constructed iDR478, a comprehensive manually curated genome-scale metabolic model (GEM) to serve as a digital twin for *E. faecium* TX0016 (strain DO). The reconstruction was based on extensive homology searches and literature evidence, and further refined and gap-filled through experimental validation. Phenotypic profiling using Biolog microarrays enabled assessment of carbon source utilization, while amino acid leave-out growth assays allowed the evaluation of auxotrophies. The final refined model accurately predicted 100% of amino acid auxotrophies and 85% of growth on sole carbon sources. Discrepancies between model predictions and experimental phenotypes identified specific knowledge gaps across metabolic pathways, including unresolved carbon source utilization phenotypes, e.g., psicose, sorbitol, and palatinose utilization. Those gaps will guide future experimental characterization. Additionally, we conducted gene essentiality analysis to evaluate the predictive capacity of iDR478. Since no experimental gene essentiality data are currently available for *E. faecium*, model predictions were compared against Tn-seq experimental results from *E. faecalis* MMH594. Under simulated unconstrained uptake conditions, iDR478 achieved 84.2% concordance with the experimental essentiality results of *E. faecalis* MMH594. iDR478 thus provides a framework for studying *E. faecium*, offers insights into its metabolic network, and serves as a source for guiding future research and identification of therapeutic targets.

## INTRODUCTION

Enterococci are Gram-positive facultative anaerobic bacteria that naturally colonize the gastrointestinal tract of humans and animals (Murray, 1990). *Enterococcus faecalis* and *Enterococcus faecium* are the most prevalent species causing nosocomial infections (Fisher & Phillips, 2009). The most common enterococcal infections include urinary tract infections (UTI), infective endocarditis, and bacteremia; they also contribute to intra-abdominal infections and meningitis (Sood et al., 2008). Among enterococci, *E. faecalis* has been the primary focus of genomic and functional studies because of its higher predominance in clinical settings (He et al., 2018; Chaguza et al., 2023); however, *E. faecium* is increasingly being recognized for its strong association with multidrug resistance (Zhou et al., 2020). As a result, *E. faecium* is now classified among the ESKAPE family of antibiotic-resistant pathogens, emphasizing the urgent need for new treatment targets (Rice, 2008).

Exploring metabolic drug targets is a promising alternative to addressing multidrug-resistant bacteria. Building metabolic network models can be a powerful tool for identifying metabolic factors linked to pathogenicity and potential antibiotic targets (Chavali et al., 2012). Despite *E. faecium*’s clinical significance, its metabolism has received limited attention, as research focused more often on antibiotic resistance mechanisms, epidemiological typing, and the characterization of virulence and colonization-associated proteins (Gao et al., 2018; Dyrkell et al., 2021; Coll et al., 2024). Consequently, aspects of *E. faecium*’s metabolism remain poorly studied, and comprehensive systems-level studies of *E. faecium* metabolism are needed.

Computational modeling and in silico simulations are potent tools that allow deep understanding of bacterial phenotypes, including metabolism. Genome-scale metabolic models (GEMs) have been valuable in elucidating genotype-phenotype relationships and generating computational predictions of biological functions (Bordbar et al., 2014). They are a key step toward *digital twins*, intended to allow full in silico simulation experiments on *virtual cells*.

GEMs are constraint-based modeling methods that translate genetic, genomic, and biochemical information into a mathematical framework for metabolic network reconstruction. They enable simulation and prediction of metabolic capabilities under various conditions (O’Brien et al., 2015). They also allow the prediction of gene essentiality by simulating the effects of multiple gene knockouts. Accordingly, GEMs have a wide range of applications, including the identification of antibiotic targets, the prediction of enzyme functions, and strain development to produce chemicals and materials (Oberhardt et al., 2009).

Thanks to a large set of well-curated models, automated metabolic reconstruction is rapidly becoming achievable (e.g., Model SEED (Devoid et al., 2013), KBase (Allen et al., 2022), and AGORA (Magnúsdóttir et al., 2017)); however, manual curation and experimental validation of GEMs for at least a reference strain per species remain valuable. Numerous GEMs have been developed for lactic acid bacteria (LAB), to which *E. faecium* belongs, which is a group characterized by significant genetic and biochemical diversity that contributes to differences in metabolic capacities and pathogenicity among species (Douillard & de Vos, 2014). Published manually curated GEMs for this group include models of *Lactococcus lactis* (Flahaut et al., 2013), *Lactiplantibacillus* (*Lactobacillus*) *plantarum* (Teusink et al., 2006), and several *Streptococcus* species, such as *S. thermophilus* (Pastink et al., 2009), *S. oralis* (Jensen et al., 2020), and *S. pyogenes* (Levering et al., 2016). Despite this progress, only one manually curated enterococcal GEM has been published for *E. faecalis* (Veith et al., 2015), leaving a significant gap in the modeling of the multidrug-resistant ESKAPE pathogen, *E. faecium*.

The rising availability of whole genome sequences has significantly accelerated the development of GEMs (Fang et al., 2020). For *E. faecium*, the first partially assembled draft genome sequence was published in 2000 for strain TX0016 (Fox, 2000), followed by the first complete genome sequence in 2012 for strain AUS004 (Lam et al., 2012) and a complete sequence for the TX0016 strain shortly thereafter (Qin et al., 2012). Today, over 33,953 genomes are available for *E. faecium*, 826 of which are described as complete (NCBI database accessed on September 30, 2025).

This study introduces iDR478, a manually curated and experimentally validated GEM for *E. faecium.* The model is reconstructed based on the genome sequence of the strain, *E. faecium* TX0016, also known as strain DO, isolated from an endocarditis patient in 1992. Our GEM enables a comprehensive exploration of the organism’s metabolic capabilities and allows for the study of its network behavior under diverse experimental conditions. We validated the model on experimental data from Biolog phenotypic arrays and amino acid auxotrophy experiments. iDR478 serves as a starting point for future enhancements and further refinements in the metabolic modeling of *E. faecium*.

## MATERIALS AND METHODS

### Strain

The strain whose genome was used in this reconstruction is *E. faecium* TX0016, also known as the DO strain (ATCC BAA-472), isolated from the blood of a patient with infective endocarditis. The DO genome sequence was downloaded from the National Center for Biotechnology Information (NCBI) (https://www.ncbi.nlm.nih.gov/, taxonomy ID: 333849) in May 2023.

### Growth conditions and media preparation

*E. faecium* TX0016 was initially cultured aerobically at 37°C in brain heart infusion (BHI) broth, as recommended by ATCC. Glycerol stocks (25%) were prepared in 2 ml cryovials and stored at -80°C for long-term preservation. For the validation experiments, the strain was grown at 37°C in a chemically defined medium for lactic acid bacteria (CDM-LAB). The CDM-LAB formulation used in our study included M9 salts, buffers, trace elements, all 20 amino acids (AAs), nucleotide bases, multiple vitamins, and diverse carbon sources. The pH of the medium was adjusted to a neutral range (6.8–7.3) by HCl or NaOH. Concentrated stock solutions of most components were prepared in advance, except for components that needed to be freshly prepared (e.g., cysteine). All stock solutions were either syringe or vacuum filter sterilized through 0.22 μm pore-size membranes and stored under appropriate storage conditions. A detailed list of medium components, preparation methods, and storage requirements is provided in **Supplementary Table S1**. The CDM-LAB composition was adapted from previously developed formulations for *E. faecium* (Zhang et al., 2013) and for the closely related *E. faecalis* (Jönsson et al., 2009; Veith et al., 2015).

### Metabolic reconstruction

To build the GEM for the *E. faecium* TX0016 strain, we used the protocol by Thiele and Palsson (2010). We began developing a metabolic network reconstruction based on genome annotation data obtained from NCBI. Candidate metabolic proteins, associated functions, potential reactions, and accession IDs were identified and curated using the information from the Bacterial and Viral Bioinformatics Resource Center, BV-BRC, formerly PATRIC (Antonopoulos et al., 2019; Olson et al., 2023) and comprehensive biochemical databases such as KEGG (Kanehisa et al., 2006), BiGG Models (King et al., 2016) and MetaNetx (Ganter et al., 2013). The curated information was compiled into a structured spreadsheet, forming the foundation of the metabolic network.

This draft reconstruction was subjected to extensive manual curation and refinement. Each metabolic reaction and function was verified against experimental data in literature, whenever possible. Because of limited specific data on *E. faecium* TX0016, literature from closely related species, most often *E. faecalis,* was used. Metabolic maps were constructed to help track and identify missing functions, using the pathway visualization tool Escher (King et al., 2015), guided by KEGG pathway maps. To fill the gaps in the network, we used BLASTp for homology searches through the NCBI web server (Boratyn et al., 2013) with default settings, relying on data from closely related species.

Each reaction or metabolite within the model was assigned a standard BiGG identifier. Metabolite identifiers were manually reviewed and standardized to ensure connectivity across the reactions in the network. For species-specific or missing pathways, not covered in BiGG databases, new metabolites and reactions were manually created and assigned unique identifiers.

Enzyme Commission (E.C.) numbers and genomic subsystem assignments were also added to all reactions, when applicable. Gene-protein-reaction (GPR) associations were assigned, linking (Boratyn et al., 2013) each reaction to its corresponding gene (via NCBI accession ID), and, when available, its gene symbol. A function may be associated with a single gene, multiple adjacent genes acting together (AND relationship), or multiple genes capable of independently performing the same function (OR relationship), depending on the genomic context.

Additionally, each reaction was also given a confidence score, reflecting the strength of the supporting available evidence on the metabolic function. The confidence scoring system (**Supplementary Table S2**) was adapted from the Thiele and Palsson’s protocol (2010), and it ranges from 0 to 4, with 0 indicating the lowest level of evidence and 4 representing the highest. The scoring and evidence data were critical for identifying low-confidence reactions that could be the reason for discordances in subsequent simulations. An additional column specifying if the function is annotated in our genome or based on homology searches was also incorporated.

Transport reactions were added into the reconstruction based on genome annotation data from NCBI, KEGG, the Transporter Classification Database, TCDB (Saier et al., 2021), and supporting literature, where available. Exchange reactions were also included to allow for metabolite uptake and secretion, completing the reconstruction network.

### Constraint-based modeling and simulations

The reconstructed metabolic network was converted into a mathematical model, with all simulations conducted using the Constraint-Based Metabolic Modeling toolbox in Python (COBRApy) version 0.25.0 (Ebrahim et al., 2013). These simulations enabled the investigation of the metabolic capabilities and adaptations through the metabolic network. Flux balance analysis (FBA) was performed using the default glpk COBRApy solver. Flux bounds were set according to the reversibility of each reaction, guided by information from the *E. faecalis* template model and the BiGG database. Stoichiometric mass and charge balance were maintained by ensuring elemental balance in each reaction.

### Gap filling

We supplemented our curated model with reactions from a CarveMe model, which constructs a model by filtering a universal (Gram-positive) model to a specific one through solving a mixed-integer linear program (Machado et al., 2018). However, the reliance on a universal model in CarveMe may limit the specificity of the model. Thus, the updated model underwent another round of intensive manual curation. Each gap-filled reaction, added by CarveMe, was thoroughly evaluated using BLASTp homology search and literature evidence, when available. A confidence score was also assigned to each added reaction to reflect the level of evidence supporting its inclusion. The model was retested for in silico biomass production. If the model remained non-functional, we performed additional gap filling using cobrapy’s default option based on Reed et al. (2006) until the model became fully functional. The gap-filled reactions were thoroughly evaluated using the same approach.

### Biomass reaction

The biomass reaction includes all the essential components required for bacterial growth, including biomass precursors, such as amino acids, nucleotide triphosphates, and lipids. Due to the lack of a biomass composition specific to *E. faecium*, we relied on the biomass reaction used in *E. faecalis* GEM (Veith et al., 2015), guided by the model Gram-positive organism, *Bacillus* (default of the CARVE-me model).

### Amino acid “leave-out” experiments

Leave-out experiments were carried out to verify the essentiality of each AA in *E. faecium* and to validate our model. Pre-cultures of *E. faecium* TX0016 cells were grown overnight at 37°C in CDM-LAB and then washed twice with phosphate buffer saline (PBS) to minimize any residual components of the rich media. The omission experiment was conducted by systematic removal of one AA from the CDM-LAB to test whether *E. faecium* can grow without it. The washed cells were inoculated into the modified media lacking the specified AA, with an initial optical density at 600 nm (OD_600_) of 0.05-0.075. The cultures were incubated overnight at 37°C in 3 ml volumes into 5 ml capped tubes. A culture grown in rich CDM-LAB with the complete AA composition was used as a positive control. Cell growth was monitored by visual inspection and by OD_600_ measurement the following day. If growth was observed, sequential inoculation steps were performed, in which cells that grew in the CDM-LAB were washed again and transferred to a fresh CDM-LAB to ensure complete removal of any residual AA from the growth medium. This step served to confirm the prototrophy of the respective AA. All omissions have been tested in at least two independently prepared media and at least three separate inoculations.

### Biolog phenotyping experiments

Biolog’s Phenotype Microarrays (Biolog, CA, USA) were used for carbon utilization assays. Kinetic measurements were read by the Omnilog^TM^ plate reader system. Carbon source utilization was tested in PM01 and PM02 plates containing 190 carbon sources. *E. faecium* cells were first cultured overnight in CDM-LAB, washed twice with PBS, and then inoculated into fresh CDM-LAB lacking glucose. This medium served as a negative control, as the strain does not grow under this condition. One hundred microliters of the inoculated cells was added into each well of the microplate and incubated at 37°C for 48 hours. Replicate experiments were conducted to ensure reproducibility and accurately determine the sole carbon utilization profiles of *E. faecium*.

### Biolog plate data processing

Kinetic time-series data were collected from the Omnilog^TM^ plate reader machine for PM01 and PM02. Each time-series signal was smoothed with a Savitzky Golay filter set to a 3rd-degree polynomial and a window size of 50. Maximum signal (respiration kinetics) values were extracted for each signal and compared to the control well maximum signal values (negative controls) with a z-test with multiple test correction to make growth calls on the carbon sources. Carbon sources with both replicates showing growth were assigned as growth-usable carbon sources, and sources with no growth across replicates were assigned as unusable carbon sources.

### Model simulations using COBRApy Amino acid auxotrophy

In silico simulations were conducted to determine auxotrophic AA(s). CDM-LAB medium was defined in silico, and model exchanges were opened up for all 20 AAs. Each AA was then removed iteratively by setting its lower and upper bounds to zero to determine the biomass yield. Growth was considered feasible if the resulting objective value exceeded a threshold of 0.01 h^-1^. If growth was not yielded upon removal of a specific AA, the organism was auxotrophic to that AA; otherwise, the organism was considered prototrophic.

### Carbon source utilization

The medium used for Biolog PM experiments (CDM-LAB without glucose) was defined in silico, and model exchanges were opened up for all carbon sources available in the Biolog phenotype microarrays. Since the model predicted in silico growth in the absence of glucose by catabolizing AAs—a phenotype not observed experimentally—a specific growth threshold was established corresponding to the in silico growth value (biomass production without added glucose). Model growth was tested on each carbon source, and only substrates generating biomass above this threshold were determined to be usable carbon sources. Discrepancies were visualized by Escher and revised against information from KEGG and BiGG. A confusion matrix was generated to express the concordance of the in silico growth outcomes with the experimental results. Experimental data on amino acid auxotrophies and carbon source utilization were incorporated to enhance the model’s accuracy.

### Gene essentiality

To assess gene essentiality, we conducted single gene knockouts in the metabolic network of *E. faecium* by subsequently performing in silico single gene deletions under two conditions: with all exchange reactions open and in defined medium. For each gene, the flux through the associated reaction was constrained to zero flux. If the resulting objective function value dropped below a threshold objective value of 0.01, the gene was classified as essential; otherwise, the gene is not essential. Gene essentiality analysis is an important measure, especially when considering new possible antimicrobial drug targets. As we could not find established experimental data on gene essentiality for the *E. faecium* TX0016 strain, we compared between iDR478 and experimental gene essentiality data using data from the closely related species, *E. faecalis* MMH594 (Gilmore et al., 2020), as a cross-species consistency assessment. Orthologs were identified between *E. faecium* DO and *E. faecalis* V583 (a strain whose locus tags are mapped to those of MMH594). The two RefSeq proteomes were compared by BLASTp, and only bidirectional best hits were retained as orthologs. Protein accession numbers were mapped to locus tags through the NCBI feature tables for each assembly. Genes assigned by Gilmore et al. (2020) as “critical, important, potentially critical, or potentially important” were counted as experimentally required for fitness (essential), and those classified as non-essential as not required (non-essential).

## RESULTS

### iDR478: a manually curated genome-scale metabolic model for *E. faecium*

The new manually curated and experimentally validated metabolic model of *E. faecium* DO (TX0016), iDR478, is based on its genome sequence (Qin et al., 2012) and includes all metabolic reactions required for cell growth. Because of scarcity of literature about *E. faecium* TX0016, we curated data from the closely related bacteria *E. faecium* AUS004 (Staerck et al., 2021), *E. faecium* E1162 (Zhang et al., 2013), and most often *E. faecalis* (Brown & Wittenberger, 1972; Hedl & Rodwell, 2004; Ramsey et al., 2014).

iDR478 comprises carbohydrate metabolism including glycolysis, the pentose phosphate pathway, and galactose, fructose, mannose, and pyruvate metabolism. It also covers nucleotide metabolism, such as purine and pyrimidine, and amino acid metabolism, including the synthesis and metabolism of all 20 AAs and the synthesis and metabolism of other peptides, such as glutathione. Additionally, it includes lipid metabolism, such as fatty acid and glycerophospholipid synthesis, as well as the synthesis of vitamins and cofactors such as folate, riboflavin, and pantothenate **(Figure 1A)**.

**Figure 1:**
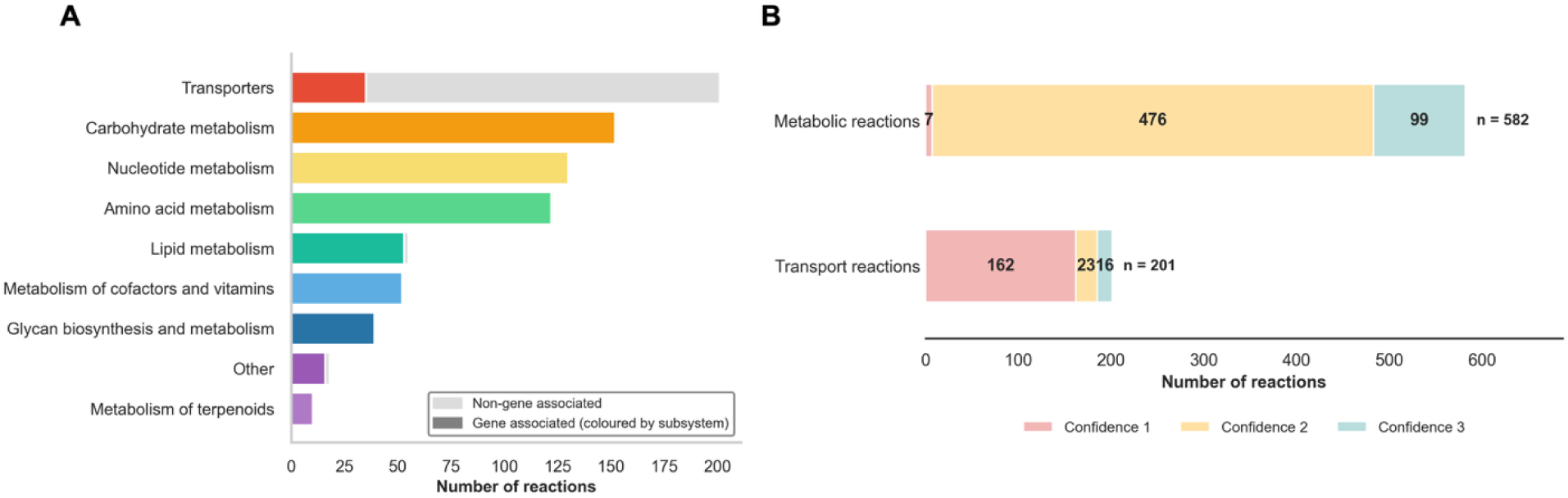
(A) Number of reactions in iDR478 in nine pathway subsystems, grouped according to KEGG metabolic pathway classification. Non-gene–associated reactions are shown in light gray at the right end of each bar. Exchange and biomass reactions are excluded. **(B)** Distribution of confidence scores across metabolic and transport reactions in iDR478. Score is shown separately for the 582 metabolic reactions and the 201 transport reactions. Exchange, boundary, and biomass reactions are excluded.

The GEM **(Table 1)** consists of 938 reactions, 638 metabolites, and 480 genes—revised to 478 genes after reactions with no connected GPRs were removed. All metabolic reactions, functions, and subsystems, as well as their confidence levels and available experimental evidence are provided in **Supplementary Table S3**. Metabolites and genes are provided as supplementary material as well **(Supplementary Tables S4 and S5, respectively)**. The model is also available in JSON and SBML formats **(Supplementary Data Sheets 1 and 2)**. Reaction confidence scores in iDR478 ranged from 1 to 3. Out of the 938 reactions, 497 (53%) were assigned a score of 2 as their inclusion evidence was only based on annotation or homology, owing to the sparsity of literature evidence available for metabolic reactions in this species **(Figure 1B)**. Eighty-four reactions were added based on homology searches following gap filling, as their associated enzymes were not initially annotated. GPRs were defined for 98.8% of the metabolic reactions (575 out of 582), excluding the biomass reaction, exchange and transport reactions. Only seven non-gene–associated reactions were included in the model because they were computed as essential to allow simulation of growth but could not be associated with a gene in the model.

**Table 1:** Characteristics of the *E. faecium* iDR478 digital twin model in terms of number of genes, reactions, and metabolites.

| Genes | Number in the model |
| --- | --- |
| Total | 478 |
| Reactions | Number in the model |
| Biomass | 1 |
| Metabolism | 582 |
| Transport | 201 |
| Exchange | 154 |
| **Total** | **938** |
| (Blocked reactions | 99)* |
| Metabolites | Number in the model |
| Intracellular | 484 |
| Extracellular | 154 |
| **Total** | **638** |

\* Blocked reactions are reactions that cannot carry any flux in simulations (Thiele & Palsson, 2010).

Many transporters in our model were not annotated as such in the reference *E. faecium* DO genome. As these had little experimental evidence available, they were mostly (162 transporters) added to support model functionality and assigned a low confidence score of 1. The final model includes 201 transport reactions that represent uptake of nutrients and export of metabolic end products, as well as 154 exchange reactions that describe the exchange of metabolites with the environment. Among the 201 transport reactions, only 16 were assigned a confidence score of 3 and 23 were assigned a score of 2 **(Figure 1B)**.

The network reconstruction provided insight into the metabolic capabilities of *E. faecium*, including the following points of interest:

Despite the limited direct experimental evidence about fatty acid biosynthesis in *E. faecium*, and given the close genetic similarities between *E. faecium* and *E. faecalis*, insights from studies on *E. faecalis* mediated by the type II fatty acid synthase (FASII) system are included in the *E. faecium model*. Fatty acids, including mainly palmitic acid (hexadecanoic acid), stearic acid (octadecanoic acid), and myristic acid (tetradecanoic acid), are attached to ACP and used as substrates in phospholipid biosynthesis (Zou et al., 2024). Glycerol-3-phosphate is initially acylated by glycerol-3-phosphate acyltransferase (PlsY, HMPREF0351_11144) then by 1-acylglycerol-3-phosphate O-acyltransferase (PlsC, HMPREF0351_12429) to form phosphatidic acid (PA). PA is subsequently converted to CDP-diacylglycerol by CDP-diacylglycerol synthetase (HMPREF0351_11681). This intermediate serves as a precursor for phosphatidylglycerol (PG) via phosphatidylglycerophosphate intermediate through phosphatidylglycerol phosphate synthase (HMPREF0351_10392), and further condensed to cardiolipin (CL) through cardiolipin synthase (HMPREF0351_1202 or HMPREF0351_11068). Experimental studies in *E. faecalis* have validated the essential roles of PlsC and acyl-ACP in this pathway, demonstrating that disruption of these enzymes leads to defective phospholipid biosynthesis and growth impairment—phenotypes that can be rescued by restoring acyl-ACP availability (Zhu et al., 2019). Fatty acid and phospholipid biosynthesis are illustrated in the Escher map **(Figure 2).**

**Figure 2:**
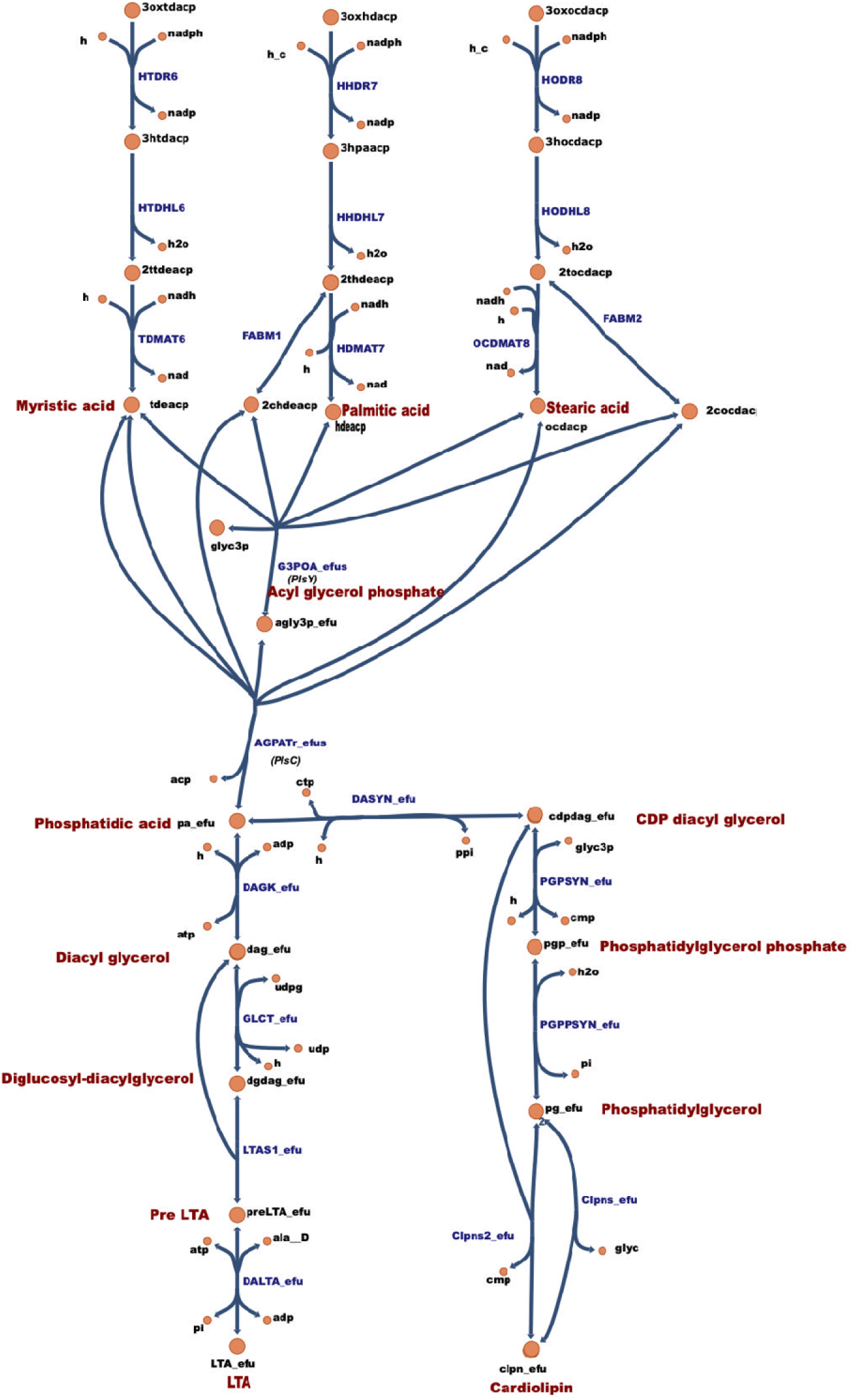
Schematic representation of a section of the fatty acid, phospholipid and lipoteichoic acid (LTA) biosynthesis pathways in *E. faecium* TX0016, as visualized by Escher. Enzymes are shown in blue, while key metabolites are highlighted in red. The pathways include the conversion of acyl-ACP to phosphatidic acid, CDP-diacylglycerol, phosphatidylglycerol, cardiolipin, and the synthesis of LTA from diacylglycerol.

*E. faecium* TX0016 also synthesizes wall teichoic acid (WTA) and lipoteichoic acid (LTA), which are critical components of its cell wall. These anionic polymers play essential roles in maintaining cell shape and division, making their representation important in GEMs for Gram-positive bacteria. The WTA biosynthesis reaction in the model reflects the experimentally defined repeating unit of *E. faecium* WTA, which consists of two N-acetyl galactosamine (GalNAc) residues linked to a glycerol-phosphate backbone (Bychowska et al., 2011; van der Es et al., 2016).These polymers are covalently attached to the peptidoglycan layer, contributing to cell wall integrity and function. Likewise, the LTA biosynthesis pathway was reconstructed based on the presence of annotated homologs of known LTA-related enzymes, enabling its incorporation into the GEM **(Figure 2)**. LTA, a key component of the Gram-positive cell envelope, consists of polyglycerol phosphate chains anchored to the cytoplasmic membrane via a glycolipid. The inclusion of the LTA biosynthetic pathway enhances the model’s ability to capture the essential features of Gram-positive envelope biosynthesis and provides a basis for exploring cell wall-targeting antimicrobials (Paganelli et al., 2017).

*E. faecium* TX0016 possesses an incomplete tricarboxylic acid (TCA) cycle, a common feature among LABs. Genomic analysis confirmed the absence of key oxidative TCA enzymes, including 2-oxoglutarate dehydrogenase and succinyl-CoA synthetase, effectively blocking the full conversion of citrate to oxaloacetate. As a result, *E. faecium* TX0016 cannot generate NADH or FADH_2_ efficiently through the TCA cycle and must rely predominantly on glycolysis and substrate-level phosphorylation for ATP production.

Homology searches exposed the absence of many enzymes in the folate biosynthesis pathway, including dihydropteroate synthase (*folP*), 2-amino-4-hydroxy-6-hydroxymethyldihydropteridine diphosphokinase (*folK*), and 7,8-dihydroneopterin aldolase (*folB*), enzymes present in *E. faecalis*, which can synthesize folate *in vivo*. Based on these findings, we hypothesize that folate is essential for *E. faecium* TX0016 and needs to be supplemented in the medium—a hypothesis that is consistent with previous studies (Ramsey et al., 2014). Furthermore, *E. faecium* TX0016 was found to be auxotrophic for eight AAs, a finding that has been supported by leave-out experiments and visualized as Escher maps.

### Biomass reaction

We relied on the biomass composition of the *E. faecalis* model, the only closely related *Enterococcus* for which a GEM was built, guided by coefficients used in the model organism of the Gram-positive bacterium *Bacillus* (default gram positive bacterium in CARVE me model). The reaction includes components essential for energy production, such as the 20 AAs, nucleotide triphosphates as ATP, lipids and essential cell wall components as LTA, WTA), cardiolipin, coenzymes as NAD, S-adenosyl-methionine, pyridoxal 5-phosphate, biotin, coenzyme A, tetrahydrofolate, thiamine diphosphate, and water. To accurately simulate the energy requirements of *E. faecium*, we incorporated the experimentally derived parameters from *E. faecalis* V583, as reported by Veith et al (2015). We added a separate ATPM reaction to simulate basal maintenance energy demand independent of biomass formation. This reaction was constrained with a lower bound of 0.81 mmol ATP/gDW/h, based on the experimentally determined mATP value reported for *E. faecalis* (Veith et al., 2015).

### Sole carbon source utilization profiles in *E. faecium* TX0016

We used the Biolog microarray system to investigate *E. faecium’*s utilization of 190 different carbon sources. For model simulations, we used the CDM-LAB medium components as the one used *in vitro* and allowed model exchanges for all carbon sources available tested in the Biolog phenotype microarrays. Out of the 190 carbon sources tested in Biolog, we studied 80 carbon sources (42.1%) that were either mapped in the metabolic model or supported growth in the Biolog experiments.

Although the CDM-LAB without glucose could not sustain growth experimentally, the model predicted a biomass yield of 0.078 h^-1^, which we used as a threshold to classify carbon sources as supporting growth or not. This discrepancy suggested the presence of false-positive growth-supporting reactions within the model. Systematic removal of individual AAs revealed that the model maintained growth when L-aspartate, L-glutamate, and L-asparagine were collectively removed. In contrast, glucose omission from this medium rendered growth infeasible which is consistent with the experimental observations. The validation outcomes of the other carbon sources were identical to those observed in the AA-rich CDM-LAB medium.

Experimentally, the strain utilized 46 carbon sources in the Biolog system. Based on these positive results, Escher maps of the corresponding pathways were generated. This allowed us to identify missing or incomplete pathways and thus reveal the gaps. Missing pathways included those for dextrin, cyclodextrins, psicose, lactulose, amygdalin, and gentiobiose. New pathways were thus reconstructed from a combination of KEGG pathway data, genome annotation, homology-based evidence and literature data. We also completed several incomplete pathways in the model, such as 5-keto-D-gluconic acid, sorbitol, and palatinose, by adding either missing reactions or transport mechanisms or both. All added reactions were incorporated in the refined final model.

Together, these additions extended the metabolic scope of the model. Before they were included, the reconstruction reproduced 49 of the 80 phenotypes (61.2%), with 21 true positives, 28 true negatives, six false positives, and 27 false negatives. Nineteen substrates changed from false negative to true positive in the refined model (**Figure 3**). After refinement, and with a selected growing threshold of 0.078 h^-1^, corresponding to the biomass yield of the model in absence of glucose, the model successfully predicted growth on 40 of the 46 Biolog-positive carbon sources, thus achieving an accuracy of 86.9%. The remaining six carbon sources, L-lyxose, D-arabinose, D-lactitol, sorbic acid, 2,3-butanedione, and L-leucine, failed to support growth in silico and thus remained false negatives, offering opportunity for discovery.

**Figure 3:**
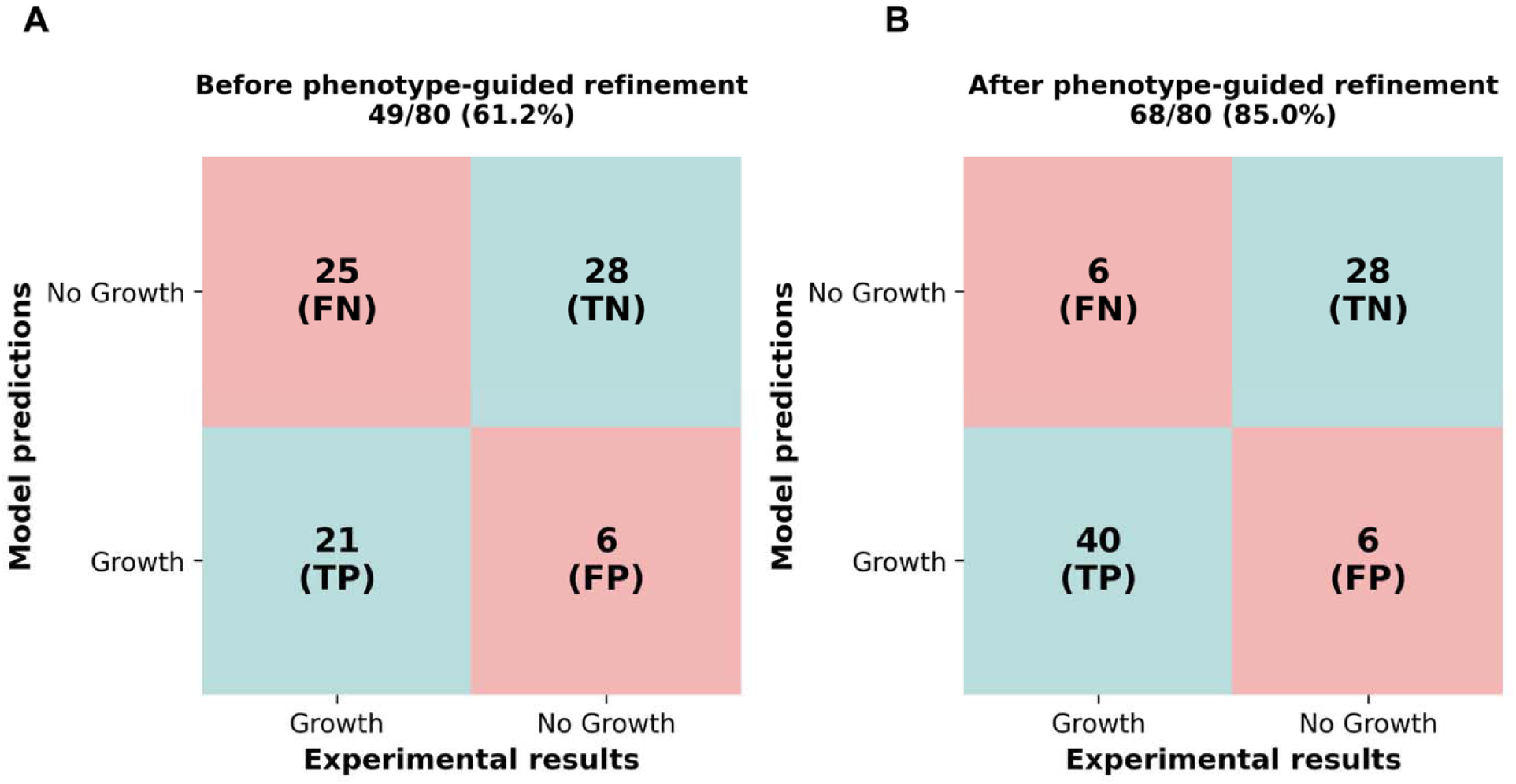
Effect of phenotype-guided refinement on predicted carbon source utilization. Confusion matrices comparing predicted growth with the Biolog phenotypes of *E. faecium* TX0016, before **(A)** and after **(B)** refinement. Agreement rose from 49 of 80 substrates (61.2%) to 68 of 80 (85%), with 21 substrates moving from false negative to true positive. **TP**: True positive, the model and the experimental data predict a positive result. **TN:** True negative, the model and experimental data predict a negative result. **FN:** False negative, the model predicts a negative result while experimental data predict a positive result. **FP:** False positive, the model predicts a positive result while experimental data predict a negative result.

Conversely, the strain did not grow on 34 mapped carbon sources, of which 28 were consistent with the model predictions, representing true negatives. However, six carbon sources, sucrose, 2’-deoxyadenosine, D-gluconic acid, thymidine, D-raffinose, and L-malic acid were false positives, as the model predicted growth that was not observed experimentally.

Overall, the agreement of the final curated iDR478 model with experimental carbon source utilization profiles was 85%, with 68 out of 80 carbon sources, 40 true positives, and 28 true negatives (**Figure 3B**). A detailed table representing the Biolog results and validations is provided in **Supplementary Table S6**.

### Amino acid essentiality in *E. faecium* TX0016

Given the limited characterization of amino acid metabolism in *E. faecium*, amino acid leave-one-out experiments were conducted to assess the amino acid essentiality and validate the newly built GEM (**Figure 4A)**. Based on experimental results, omission of L-methionine, L-histidine, L-leucine, L-isoleucine, L-valine, L-arginine, L-threonine, and L-tryptophan resulted in a marked decrease in the growth, with final OD_600_ values consistently below 0.1 across all replicates. Thus, these eight AAs are classified as essential for growth. The strain initially failed to grow in the absence of L-lysine, L-phenylalanine, and L-tyrosine. However, after repeated re-inoculation, the cultures adapted and demonstrated significantly improved growth, indicating a potential for adaptation upon passaging **(Figure 4B**). An OD_600_ above 0.21 was observed across all experiments in cultures where L-alanine, L-proline, L-glutamate, L-glutamine, L-aspartate, and L-asparagine were omitted. The essentiality of L-serine, L-glycine, and L-cysteine remained inconclusive, as their omission resulted in a weak growth (OD_600_ valued above 0.1 but below 0.21).

**Figure 4:**
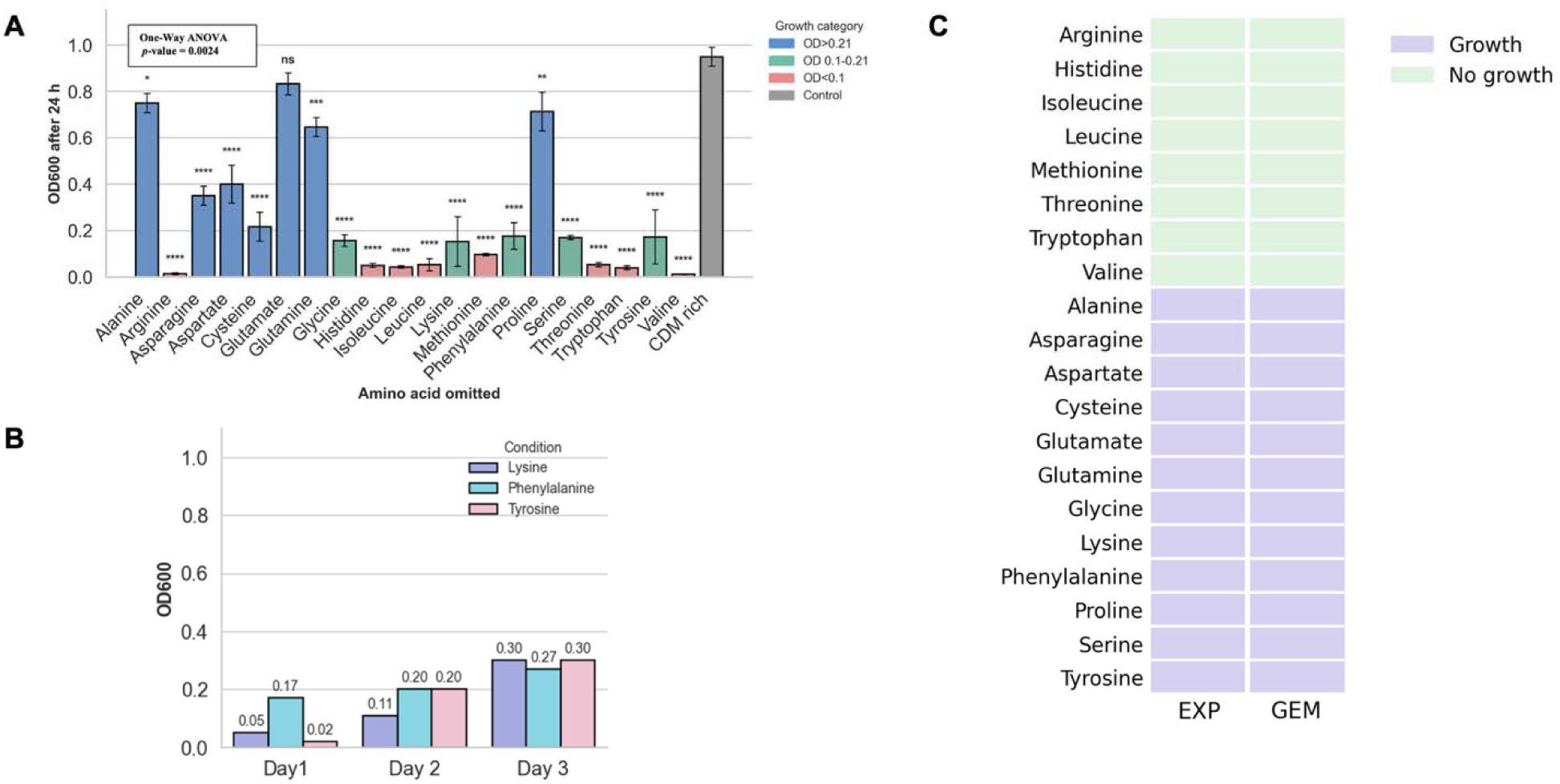
Amino acid auxotrophy experiments of *E. faecium* DO. **(A)** Mean final optical density (OD_600_) of *E. faecium* DO in the absence of single amino acids from the CDM-LAB. Bar chart showing final mean OD_600_ values after 24 hours incubation. Each bar represents the mean OD of three biological replicates ± SD for a condition where single amino acid was omitted. Cultures with a final OD < 0.1 are marked in red; those with a final OD > 0.21 are marked in blue; and those with weak growth (OD between 0.1 and 0.21) in green. A one-way ANOVA test was performed to compare growth (final average OD_600_) across amino acid omissions. Results were significantly different (*p* value = 0.0024, ≤ 0.05). This was followed by Tukey’s multiple comparisons *post hoc* test. Asterisks indicate adjusted *p*-values corresponding to multiple comparisons against the complete CDM-LAB control: \**p* < 0.05, \*\**p* < 0.01, \*\*\**p* < 0.001, \*\*\*\**p* < 0.0001; ns = not significant. **(B)** Repeated passaging of cultures grown in the absence of lysine, phenylalanine, and tyrosine, respectively, results in adaptation to omissions. **(C)** Individual comparison of each amino acid between the experimental results of the amino acid leave-out experiments (EXP) and the simulation results of the model iDR478 (GEM).

With an OD_600_ threshold of 0.1 selected based on a sensitivity analysis presented (**Supplementary Table S7**), the model achieved 100% accuracy in predicting amino acid essentiality, identifying eight essential AAs, L-methionine, L-histidine, L-leucine, L-isoleucine, L-valine, L-arginine, L-threonine, and L-tryptophan and 12 non-essential AAs, L-alanine, L-proline, L-glutamate, L-glutamine, L-aspartate, L-asparagine, L-lysine, L-phenylalanine, and L-tyrosine, L-serine, L-glycine, and L-cysteine **(Figure 4C)**.

### Gene essentiality

Gene essentiality was assessed under two conditions: (i) with all exchange reactions open (simulating rich media) and (ii) in CDM.

With all the exchange reactions open, the model predicted 19 essential genes, including genes from nucleotide metabolism (n = 4) and amino acid metabolism (n = 1). The largest proportion of essential genes, however, was assigned to the metabolism of cofactors and vitamins (n = 12), such as pantothenate and CoA biosynthesis and nicotinate and nicotinamide metabolism. In CDM, 88 essential genes were predicted in the model, including genes from nucleotide metabolism (n = 17), lipid metabolism (n = 12), amino acid metabolism (n = 18), peptidoglycan and glycan biosynthesis (n = 16), metabolism of cofactors and vitamins (n = 13), and the terpenoid backbone pathway (n = 7). All 19 genes that are essential under unconstrained uptake are also essential in CDM-LAB (**Figure 5A)**.

**Figure 5:**
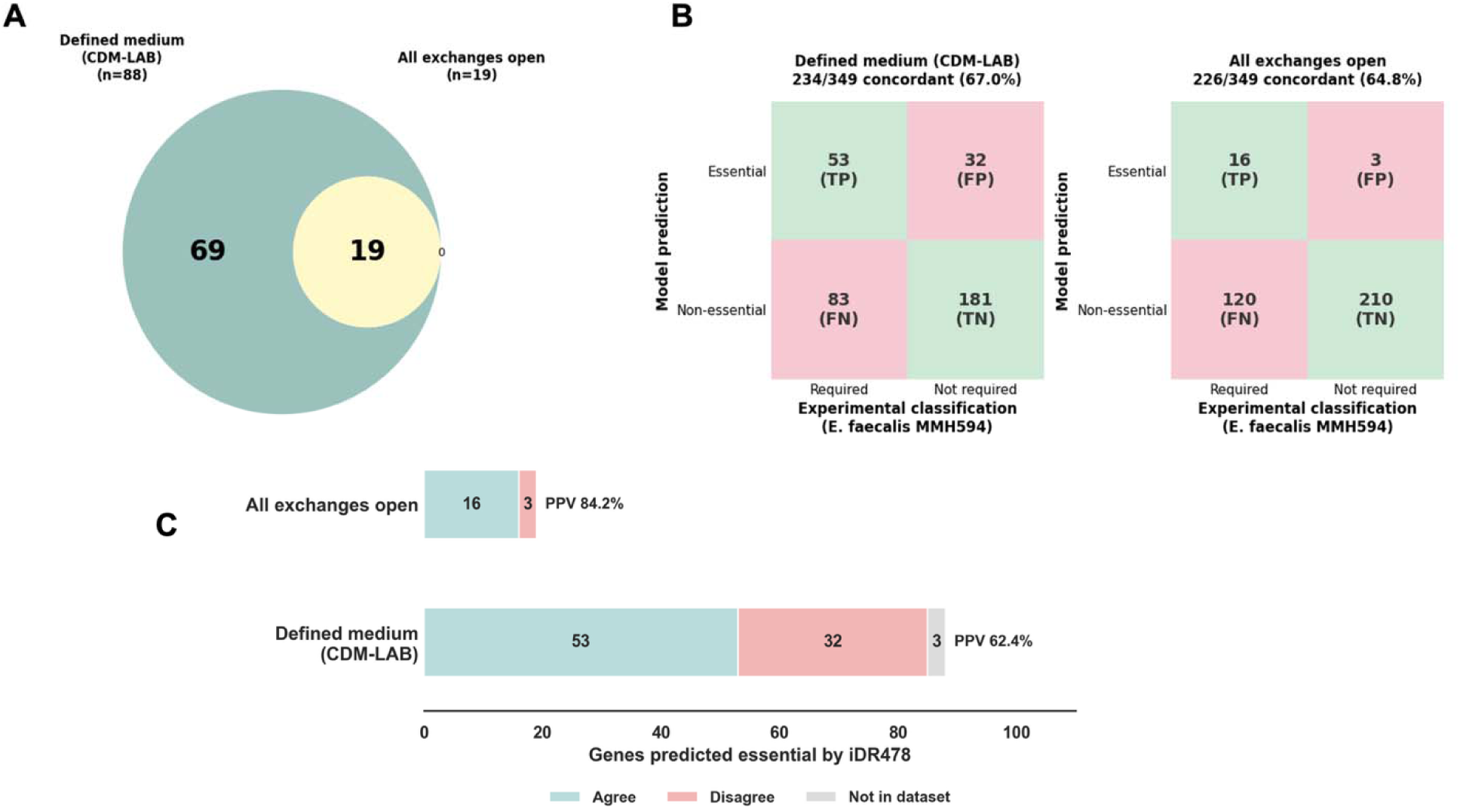
(A) Venn diagram illustrating the shared/unique essential genes between the two conditions (a chemically defined medium and one with all exchanges open). **(B)** Confusion matrices comparing the predictions with the fitness classification of orthologous genes in *E. faecalis* MMH594 (Gilmore et al., 2020), across the 349 genes of iDR478 for which a reciprocal best-hit ortholog with a fitness classification was available. **TP:** predicted essential and experimentally required; **TN:** predicted non-essential and not required; **FP:** predicted essential but not required; **FN:** predicted non-essential but required. **(C)** Comparison of the genes predicted to be essential under each condition (positive predicted value, PPV), showing those whose classification agrees with the experimental assignment, those that disagree, and those for which no ortholog with a fitness classification was available.

Since no experimental essentiality data were available for *E. faecium* TX0016, the model’s predictions were compared with a Tn-seq study of MDR *E. faecalis* MMH594 (Gilmore et al., 2020).

Orthologs were identified by reciprocal best BLASTp hits between the *E. faecium* DO and *E. faecalis* and 352 of the 478 genes of the model have an ortholog, and 349 of these carry a fitness classification in Gimlore’s study. Genes classified in the experimental data as critical, important, potentially critical, or potentially important are counted as required for fitness, and those classified as non-essential as not required. The analysis was repeated, counting only the critical and important categories and excluding potentially important categories, under which agreement is marginally higher (Matthews correlation coefficient, MCC, of 0.31 *vs.* 0.27 in defined medium), so the inclusive mapping reported here is the more conservative.

In the defined medium, 85 of the 88 predicted essential genes have an ortholog in the *E. faecalis* experimental dataset. Among these, 53 genes (62.4%) agreed with the experimental classification, giving true positive results. Across the 349 assessable genes, the whole comparison indicates 181 true negatives, 32 false positives, and 83 false negatives, corresponding to a sensitivity of 0.39, a specificity of 0.85, a positive predictive value (PPV) of 0.624, an accuracy of 0.67, a balanced accuracy of 0.62, and an MCC of 0.27. Under unconstrained uptake, 16 of the 19 predicted essential genes (84.2%) agreed with the experimental classification, with 210 true negatives, three false positives, and 120 false negatives, a PPV of 0.842, a specificity of 0.986 but accuracy of 0.648, a balanced accuracy of 0.552, and an MCC of 0.223 **(Figures 5B and 5C)**. The full comparison, including the ortholog assignment and the experimental classification for every gene, is provided in **Supplementary Table S8**.

The model also predicted 31 essential reactions, when all the exchanges are open, including reactions from the pentose phosphate pathway (n = 1, e.g., phosphoribosylpyrophosphate synthetase), nucleotide metabolism (n = 2, e.g., UMP kinase), and cofactor and vitamin metabolism (n = 9), such as pantothenate and CoA biosynthesis and vitamin B6 metabolism pathways. It also included transporters (n = 3) and exchange reactions (n = 9). In the CDM, 140 essential reactions were predicted, including 15 exchanges (eight essential AAs and six cofactors) and their transporters (n = 6), as well as reactions from carbohydrate metabolism (n = 4), amino acid metabolism (n= 14), metabolism of cofactors and vitamins (n = 11), nucleotide metabolism (n = 16), and the terpenoid backbone pathway (n = 8) with the largest proportion of essential reactions assigned to lipid metabolism (n = 43), followed by cell wall and glycan biosynthesis (n = 19). A complete list of predicted essential reactions is provided in **Supplementary Table S9**.

## DISCUSSION

We here present iDR478, a comprehensive manually curated genome-scale metabolic model of *E. faecium* DO (TX0016), a major multidrug-resistant member of the ESKAPE pathogens. The model serves as an established platform for conducting systems-level analyses of an organism’s metabolism by integrating data from genome annotation, biochemical characterizations, and literature. Model curation integrated existing knowledge of *E. faecium* annotations and metabolic pathways with insights from our experimental analyses Reconstruction was guided and refined through experimental validation, using Biolog phenotypic microarrays for carbon source utilization and amino acid leave-out growth assays for amino acid auxotrophies. After refinement, the model fully predicted amino acid auxotrophies in agreement with leave-out epxeriments, and achieved 85% agreement with Biolog’s sole carbon source utilization.

The metabolic potential of *E. faecium* was evaluated across 80 carbon sources. Model validation for carbon source utilization was conducted via growth simulation in a CDM-LAB containing all AAs but lacking glucose, which exactly mirrors the conditions of experimental Biolog carbon utilization assays. The model’s ability to sustain growth in glucose-free conditions, despite the absence of experimental growth, was likely driven by false-positive utilization of L-aspartate, L-glutamate, and L-asparagine. Removing all three together with glucose abolished predicted growth, whereas removing any two did not. This finding suggests that the model catabolizes these AAs for energy generation, leading to false-positive growth predictions under the Biolog-like conditions. Mapping and pathways analysis revealed that these AAs are interconverted through transamination and deamidation reactions, ultimately supplying carbon to the oxaloacetate node and allowing the model to synthesize essential biomass precursors. This behavior is unlikely to occur under the experimental conditions, suggesting that the current reconstruction overestimates the capacity of these AAs pathways to sustain growth.

Out of the 80 carbon sources tested, 40 were predicted by the model as true positives and 28 as true negatives. However, the predictions of six substrates were false negative, as they failed to support carbon utilization in the model although the substrates were effectively utilized experimentally. These discrepancies likely result from gaps in our understanding of specific metabolic pathways. For instance, sorbic acid and D-lactitol utilization failed in the model because of incomplete knowledge of the metabolic mechanisms of these substrates, as recovered from literature searches. As for D-arabinose, it could not be utilized in silico because of the absence of arabinose isomerase in the genome, with neither literature nor genomic evidence supporting its presence. Similarly, L-lyxose and 2,3-butanedione (diacetyl) metabolism remain unsupported owing to missing key enzymes, such as ATP: L-xylulose 5-phosphotransferase and diacetyl reductase, respectively. There is neither genomic evidence nor literature support for the presence of these enzymes or any homolog thereof in any *Enterococcus* species. Another false negative result is L-leucine, transaminated to 4-methyl-2-oxopentanoate, which is usually converted into isovaleryl-CoA through an intermediate step, but this needs a specific enzyme complex (BCKDH) that also could not be found in the genome sequence.

Conversely, six carbon sources represented false positives predictions of the model, as they were predicted to support growth in silico but were not experimentally validated. Sucrose was predicted as a utilized carbon source in the model based on two annotated sucrase enzyme-encoding genes in the model (HMPREF0351_11791 or HMPREF0351_11800, EC: 3.2.1.26, BiGG ID SUCR), yet no respiration was observed in Biolog experiments. Literature evidence also confirmed the presence of universal sugar transport system components in *E. faecium* (Hallenbeck et al., 2024). Thus, this discordance may be due to inducible expression requirements that Biolog conditions did not fulfill, since sucrase has previously been studied as an inducible enzyme in Gram-positive bacteria, e.g., in a previous *Staphylococcus* study (Gering & Brückner, 1996). Likewise, raffinose, thymidine, and deoxydenosine utilization in the model was based on nonspecific broad-spectrum enzymes, notably alpha-galactosidase (HMPREF0351_11597, EC: 3.2.1.22, BiGG ID: RAFGH) in the case of raffinose; pyrimidine-nucleoside phosphorylase (HMPREF0351_10038, EC: 2.4.2.4, BiGG ID: TMDPP) in the case of thymidine; and purine-nucleoside phosphorylase (PNP) (HMPREF0351_10046, EC: 2.4.2.1, BiGG ID: PUNP2) in the case of deoxyadenosine. These observations emphasize how annotation-based assumptions may overestimate metabolic capabilities. In the case of deoxyadenosine, PNP may possibly prefer to act on other nucleosides such as inosine (which was indeed utilized both in silico and experimentally). The inability of raffinose to support the strain growth in real life may be also interpretable by the non-functionality of the enzyme in all *E. faecium* strains, as studied before. This capability was shown to be linked to a specific gene cluster located on a megaplasmid, encoding two alpha galactosidase enzymes, AgaA and AgaB, present uniquely in some strains and not in others (Zhang et al., 2011). Similar issues were noted in gluconate and L-malic acid utilization. Gluconate utilization was also predicted by the model but not observed experimentally. The model includes key enzymes for its catabolism: gluconokinase (HMPREF0351_11707, EC 2.7.1.12, BiGG ID: GLUKA_1) and 6-phosphogluconate dehydrogenase (EC 1.1.1.44). A likely explanation is the absence of a gluconate transporter in *E. faecium* TX0016, as the gluconate transporter included in our model was supported by a low-confidence score of 1 (added primarily just to ensure model connectivity).

For L-malic acid, misprediction may have stemmed from incorrect assumptions regarding maltose-related enzymes. The model’s prediction relies on the annotated maltose phosphorylase enzyme (HMPREF0351_10418, EC: 2.4.1.8, BiGG ID: MPL), which is assigned a high confidence score of 3. This score is supported by both genome annotation and literature evidence from *E. faecalis* (Mokhtari et al., 2013), which describes maltose catabolism via maltose phosphorylase (MalP) and MapP enzymes. Additionally, the presence of a PEP-dependent maltose PTS transporter in the model is also supported by universal PTS system (PtsIH) proteins in *E. faecium* (Hallenbeck et al., 2024) and *E. faecalis* (Mokhtari et al., 2013). While these studies suggest a potential for maltose-related metabolism, they may have led the model to incorrectly assume that it supports growth on L-malic acid. The lack of positive Biolog results suggests that key steps are inactive under tested conditions or require regulatory induction.

Alternatively, adaptation or multiple passages might be necessary to activate the pathway, highlighting the difference between genomic potential and observable phenotype. Similar considerations can be applied to all the previously mentioned false positive cases, where transporters or relevant metabolic enzymes may be conditionally expressed or not induced under Biolog assay conditions. Thus, it is possible that, with further passaging, the strain might eventually activate the required pathways for alternative carbon source utilization.

The difference between genomic potential and observable phenotype in all false positive cases highlights a common limitation of GEMs. GEM predictions generally assume that all genetically encoded reactions are available, whereas *in vivo* enzyme activity is strictly regulated by gene expression and environmental context. Thus, incorporating transcriptomic and regulatory data could significantly improve predictive accuracy (Rychel et al., 2021; Catoiu et al., 2025).

The metabolic network was further validated against experimentally determined amino acid auxotrophies. Amino acid leave-out experiments demonstrated strong agreement between in silico predictions and experimental outcomes. Eight AAs, L-methionine, L-histidine, L-leucine, L-isoleucine, L-valine, L-arginine, L-threonine, and L-tryptophan, were deemed essential, as the average optical density for cultures lacking them remained below 0.1 across all experiments after 24 hours. These findings align with the model predictions as true negative results. It is worth noting, however, that these amino acid-negative conditions were not subjected to repeated passaging. Therefore, the possibility of adaptation over time cannot be excluded. In contrast, the remaining 12 AAs, L-alanine, L-proline, L-glutamate, L-glutamine, L-aspartate, L-asparagine, L-lysine, L-phenylalanine, and L-tyrosine, L-serine, L-glycine, and L-cysteine were considered as nonessential, merely following the selected threshold of growth and the model predicted prototrophy.

Digging deeper in the experimental results, beyond just the sensitivity analysis approach and the selected threshold, we further analyzed the exact OD_600_ values for each condition and assessed the growth patterns across experiments and biological triplicates. The six AAs L-alanine, L-proline, L-glutamate, L-glutamine, L-aspartate, and L-asparagine are surely classified as non-essential for growth, as cultures consistently exhibited OD_600_ values above 0.21 across all experiments, and the model also predicted prototrophy, representing true positive results.

However, upon initial omission of any of L-lysine, L-phenylalanine, and L-tyrosine, the strain grew marginally. Interestingly, following repeated passaging, the final optical density was well above the threshold, suggesting that a regulatory or metabolic adaptation may have occurred during passaging. These cases ultimately align with model predictions of prototrophy.

Experimental growth in the absence of L-serine, L-glycine, and L-cysteine was consistently marginal, and the model predicted nonessentiality. The uncertainty surrounding L-glycine and L-serine essentiality may be linked to the metabolic constraints of glycine hydroxymethyltransferase (HMPREF0351_12288, BiGG ID: GHMT2r, *glyA*), which catalyzes the reversible conversion of L-serine and tetrahydrofolate (THF) to glycine and 5,10-methylenetetrahydrofolate (MTHF). Since *E. faecium* cannot synthesize folate *de novo* (Ramsey et al., 2014) and the CDM medium contains only traces of folate, limited folate availability may impair GHMT2r activity, restricting growth. The model does not account for kinetic constraints or real-world nutrient limitations, which may contribute to this discrepancy. The other source of glycine in the model is aminopeptidase, which is a broad-spectrum nonspecific enzyme with no direct evidence of its presence to produce glycine specifically. Key enzymes in the phosphoglycerate pathway of serine biosynthesis, the enzyme phosphoserine aminotransferase (EC: 2.6.1.52) that converts 3-phosphohydroxypyruvate to 3-phosphoserine and the enzyme phosphoserine phosphatase (EC: 3.1.3.3) that converts 3-phosphoserine to L-serine, are also lacking, which also confirms that L-serine production primarily relies on the GHMT2r reaction as well.

Similar observations were made in GEM of *E. faecalis* V583, although their explanation evolved around transcriptomic regulation rather than folate insufficiency, given its ability to synthesize folate (Veith et al., 2015). In *Lactococcus lactis*, *the glyA* gene that codes for GHMT, and other folate metabolism genes is regulated by the PurR repressor, whose activity is influenced by hypoxanthine and PRPP. A PurR repressor protein was also identified in the genome of *E. faecalis* (purR, EF0058), displaying notable sequence similarity to homologs in *Bacillus subtilis* (55% identity) and *Lactococcus lactis* MG1363 (53% identity). Furthermore, a PurR binding motif closely resembling the consensus sequences characterized in *B. subtilis* and *L. lactis* was detected upstream of the *glyA* gene in *E. faecalis*, implying a potentially conserved PurR-dependent regulatory mechanism within this species.

For L-cysteine, the observed growth in its absence was also minimally above the threshold, with an average OD_600_ of approximately 0.25. One possible explanation is that cysteine biosynthesis requires hydrogen sulfide (H_2_S) as a sulfur donor for the cysteine synthase reaction (HMPREF0351_10472 or HMPREF0351_11313, EC: 2.5.1.47, BiGG ID: CYSS).

However, only trace amounts of H_2_S are present under the ambient air conditions used in the experiments, conditions that differ from the ideal assumptions of the in silico model (there is an exchange reaction for H_2_S as in *E. faecalis* GEM too). Furthermore, this strain lacks key genes involved in the reduction of sulfate to H_2_S, specifically *sat* (sulfate adenylyltransferase) and *cysH* (phosphoadenosine phosphosulfate reductase). These genetic limitations, combined with the low environmental H_2_S availability, likely explain the limited growth observed in the absence of L-cysteine and the discordance from the predictions of the in silico model. These findings highlight the need for integrating more kinetic constraints into GEM simulations.

Additionally, essential gene predictions were performed in the model under two simulated conditions: rich medium with unrestricted exchanges and in CDM-defined growth. Because of the lack of experimental essentiality data for *E. faecium*, we used Tn-seq data from *E. faecalis* (Gilmore et al., 2020) for cross-species consistency assessment. As a result, 84.2% of the predicted essential genes in case of the unrestricted exchanges and 62.4% in case of the CDM growth aligned with experimentally determined essential genes, representing the PPV. A full confusion matrix represents moderate agreement between the species regarding essentiality. These differences likely reflect high species-specific variations between *Enterococcus* species, differences in growth conditions, as well as limitations of the model as the absence of regulatory and adaptive constraints. This highlights the need for generating *E. faecium* specific experimental gene essentiality datasets to improve model accuracy in this aspect. Experimental approaches such as targeted gene knockout or CRISPRi studies, as demonstrated in previous studies for different genes including *acpH, treA*, and *lacL* (de Maat et al., 2019; Chua & Collins, 2022) will be critical in validating gene essentiality and identifying potential antibiotic targets in the future.

It should be noted that the quality and completeness of a metabolic reconstruction are highly dependent on the availability of accurate organism-specific data. For *E. faecium*, this remains a significant limitation, as the organism is much less characterized than related species, and comprehensive metabolic experimental datasets are not widely available. This limitation is especially critical for *E. faecium*, given its multidrug resistance; in such case, accurate metabolic models could be important tools to identify essential pathways and thus potential therapeutic targets.

iDR478 represents a manually curated GEM that provides a foundation for understanding *E. faecium’*s metabolism and predicting nutrient dependencies and essential pathways, which can guide targeted Tn-seq or CRISPR-based screens and thus inform the design of targeted antimicrobial strategies in future studies.

## DECLARATIONS

### Dedication

The authors dedicate this manuscript to the soul of the co-author, Yomna A. Hashem^¶^, who was deceased on March 30^th^, 2026, before seeing this manuscript submitted. The first complete version of the manuscript was submitted as a preprint on what would have been her 40^th^ birthday, May 1^st^, 2026.

### Author contributions

Conceptualization: BOP, JMM, RKA; methodology: JK, BOP, JMM; software: DSR, JK, JMM; validation: DSR, JK, YAH, MTK, RKA; formal analysis: DSR, JK, YAH, MTK; investigation: DSR, JK; resources: YAH, BOP, MTK, JMM; data curation; DSR, JK; writing–original draft: DSR, JK, MTK, JMM; writing–review and editing: MTK, JMM, RKA; visualization: DSR, JK, JMM; supervision YAH, MTK, JMM, RKA; all authors checked the different drafts and all (except YAH^¶^) approved the final submitted version.

## Acknowledgment and funding

The work described here was not specifically funded by a research grant. Dina Rasmi acknowledges the U.S. Agency for International Development (USAID) program, through which she received a short-term scholarship in 2023 that allowed her to visit the laboratory of Jonathan Monk, PhD and Bernhard Palsson, PhD. The scholarship was part of the “ninth announcement of the Higher Education Initiative - Graduate Scholarships for Professionals (GSP).” The funder had no interference with the content of this manuscript or the decision to publish it. The scholarship did not include a budget to cover publication fees.

## Competing interests

The authors have no personal or commercial competing interests to declare regarding the publication of this manuscript. No professional medical writing assistance was used.

## Ethical committee approval

No ethical approval was required for this work, as it does not involve human subjects or experimental animals.

## Use of generative AI

The authors have not used large language models to generate any of the text in this manuscript.

## SUPPLEMENTARY MATERIAL

**Supplementary Table S1:** A detailed list of the CDM-LAB medium components, preparation methods, and storage requirements.

**Supplementary Table S2:** Confidence scoring system, adapted from Thiele and Palsson’s protocol (2010).

**Supplementary Table S3:** Reactions in iDR478 *E. faecium* model

**Supplementary Table S4:** Metabolites in the iDR478 *E. faecium* model

**Supplementary Table S5:** Genes in the iDR478 *E. faecium* model

**Supplementary Table S6:** Carbon source utilization comparison between Biolog data and iDR478 model

**Supplementary Table S7:** Sensitivity analysis conducted to select the growth threshold in amino acid auxotrophy experiments.

**Supplementary Table S8:** A list of genes assessed for essentiality in iDR478 and their concordance with experimental data

**Supplementary Table S9:** A list of the predicted essential reactions in iDR478

**Data Sheet S1:**iDR478 model in Json format

**Data Sheet S2**: iDR478 model in SBML format

## Notes

### Competing Interest Statement

The authors have declared no competing interest.

### Summary of Updates

This third version (v3) of the manuscript has been revised, after a few months of the first submission, to include the following updates: - The title has been changed to be more descriptive and to accurately reflect the distinctiveness of the model: that it is manually curated and validated by two sets of high-throughput experiments - The model itself has been revised and its reactions and genes were updated. The model name now is iDR478 instead of iDR479 to reflect the revised number of genes. - As we received feedback about certain aspects of the manuscript content, we updated most of the figures (Figures 1, 3, 4, and 5 were updated) and thus their legends, and we also included some details on the cross-species essentiality comparisons - We updated several values related to the validation, including for example the positive predictive value (PPV) and the %agreement between essentiality studies. - We used more accurate terms to describe the Biolog screens (e.g., referring to carbon-source utilization instead of simply growth/no growth phenotypes) - Based on feedback during post-submission peer review, we added three references, including one about the AGORA (Assembly of Gut Organisms through Reconstruction and Analysis) resource, which includes semi-automated models of gut bacteria, including some E. faecalis strains - Based on repeated feedback, we avoided describing our current reconstruction as a 'metabolic digital twin'. We believe it is rather a serious step toward a digital twin for this ESKAPE pathogen.

## REFERENCES

Allen, B. H., Gupta, N., Edirisinghe, J. N., Faria, J. P., & Henry, C. S. (2022). Application of the metabolic modeling pipeline in KBase to categorize reactions, predict essential genes, and predict pathways in an isolate genome. Methods Mol. Biol., 2349, 291–320. 10.1007/978-1-0716-1585-0_13

Antonopoulos, D. A., Assaf, R., Aziz, R. K., Brettin, T., Bun, C., Conrad, N., … Yoo, H. (2019). PATRIC as a unique resource for studying antimicrobial resistance. Brief. Bioinform., 20(4), 1094–1102. 10.1093/bib/bbx083

Boratyn, G. M., Camacho, C., Cooper, P. S., Coulouris, G., Fong, A., Ma, N., … Zaretskaya, I. (2013). BLAST: a more efficient report with usability improvements. Nucleic Acids Res., 41(Web Server issue), W29-33. 10.1093/nar/gkt282

Bordbar, A., Monk, J. M., King, Z. A., & Palsson, B. O. (2014). Constraint-based models predict metabolic and associated cellular functions. Nat. Rev. Genet., 15(2), 107–120. 10.1038/nrg3643

Brown, A. T., & Wittenberger, C. L. (1972). Induction and regulation of a nicotinamide adenine dinucleotide-specific 6-phosphogluconate dehydrogenase in *Streptococcus faecalis*. J. Bacteriol., 109(1), 106–115. 10.1128/jb.109.1.106-115.1972

Bychowska, A., Theilacker, C., Czerwicka, M., Marszewska, K., Huebner, J., Holst, O., … Kaczyński, Z. (2011). Chemical structure of wall teichoic acid isolated from *Enterococcus faecium* strain U0317. Carbohydr. Res., 346(17), 2816–2819. 10.1016/j.carres.2011.09.026

Catoiu, E. A., Krishnan, J., Li, G., Lou, X. A., Rychel, K., Yuan, Y., … Palsson, B. O. (2025). iModulonDB 2.0: dynamic tools to facilitate knowledge-mining and user-enabled analyses of curated transcriptomic datasets. Nucleic Acids Res., 53(D1), D99–d106. 10.1093/nar/gkae1009

Chaguza, C., Pöntinen, A. K., Top, J., Arredondo-Alonso, S., Freitas, A. R., Novais, C., … Corander, J. (2023). The population-level impact of *Enterococcus faecalis* genetics on intestinal colonization and extraintestinal infection. Microbiol. Spectr., 11(6), e0020123. 10.1128/spectrum.00201-23

Chavali, A. K., D’Auria, K. M., Hewlett, E. L., Pearson, R. D., & Papin, J. A. (2012). A metabolic network approach for the identification and prioritization of antimicrobial drug targets. Trends Microbiol., 20(3), 113–123. 10.1016/j.tim.2011.12.004

Chua, M. J., & Collins, J. (2022). Rapid, efficient, and cost-effective gene editing of *Enterococcus faecium* with CRISPR-Cas12a. Microbiol. Spectr., 10(1), e0242721. 10.1128/spectrum.02427-21

Coll, F., Gouliouris, T., Blane, B., Yeats, C. A., Raven, K. E., Ludden, C., … Peacock, S. J. (2024). Antibiotic resistance determination using *Enterococcus faecium* whole-genome sequences: a diagnostic accuracy study using genotypic and phenotypic data. Lancet Microbe, 5(2), e151–e163. 10.1016/s2666-5247(23)00297-5

de Maat, V., Stege, P. B., Dedden, M., Hamer, M., van Pijkeren, J. P., Willems, R. J. L., & van Schaik, W. (2019). CRISPR-Cas9-mediated genome editing in vancomycin-resistant *Enterococcus faecium*. FEMS Microbiol. Lett., 366(22). 10.1093/femsle/fnz256

Devoid, S., Overbeek, R., DeJongh, M., Vonstein, V., Best, A. A., & Henry, C. (2013). Automated genome annotation and metabolic model reconstruction in the SEED and Model SEED. Methods Moll. Biol., 985, 17–45. 10.1007/978-1-62703-299-5_2

Douillard, F. P., & de Vos, W. M. (2014). Functional genomics of lactic acid bacteria: from food to health. Microb. Cell Fact., 13 Suppl 1(Suppl 1), S8. 10.1186/1475-2859-13-s1-s8

Dyrkell, F., Giske, C. G., & Fang, H. (2021). Epidemiological typing of ST80 vancomycin-resistant *Enterococcus faecium*: core genome multilocus sequence typing versus single nucleotide polymorphism-based typing. J. Glob. Antimicrob. Resist., 25, 119–123. 10.1016/j.jgar.2021.03.005

Ebrahim, A., Lerman, J. A., Palsson, B. O., & Hyduke, D. R. (2013). COBRApy: COnstraints-Based Reconstruction and Analysis for Python. BMC Syst. Biol., 7, 74. 10.1186/1752-0509-7-74

Fang, X., Lloyd, C. J., & Palsson, B. O. (2020). Reconstructing organisms in silico: genome-scale models and their emerging applications. Nat. Rev. Microbiol., 18(12), 731–743. 10.1038/s41579-020-00440-4

Fisher, K., & Phillips, C. (2009). The ecology, epidemiology and virulence of *Enterococcus*. Microbiology (Reading), 155(Pt 6), 1749–1757. 10.1099/mic.0.026385-0

Flahaut, N. A., Wiersma, A., van de Bunt, B., Martens, D. E., Schaap, P. J., Sijtsma, L., … de Vos, W. M. (2013). Genome-scale metabolic model for *Lactococcus lactis* MG1363 and its application to the analysis of flavor formation. Appl. Microbiol. Biotechnol., 97(19), 8729–8739. 10.1007/s00253-013-5140-2

Fox, J. (2000). DOE Joint Genome Institute Feat: one-day sequencing of *E. faecium*. ASM News, 66(7).

Ganter, M., Bernard, T., Moretti, S., Stelling, J., & Pagni, M. (2013). MetaNetX.org: a website and repository for accessing, analysing and manipulating metabolic networks. Bioinformatics, 29(6), 815–816. 10.1093/bioinformatics/btt036

Gao, W., Howden, B. P., & Stinear, T. P. (2018). Evolution of virulence in *Enterococcus faecium*, a hospital-adapted opportunistic pathogen. Curr. Opin. Microbiol., 41, 76–82. 10.1016/j.mib.2017.11.030

Gering, M., & Brückner, R. (1996). Transcriptional regulation of the sucrase gene of *Staphylococcus xylosus* by the repressor ScrR. J. Bacteriol., 178(2), 462–469. 10.1128/jb.178.2.462-469.1996

Gilmore, M. S., Salamzade, R., Selleck, E., Bryan, N., Mello, S. S., Manson, A. L., & Earl, A. M. (2020). Genes contributing to the unique biology and intrinsic antibiotic resistance of *Enterococcus faecalis*. mBio, 11(6). 10.1128/mBio.02962-20

Hallenbeck, M., Chua, M., & Collins, J. (2024). The role of the universal sugar transport system components PtsI (EI) and PtsH (HPr) in *Enterococcus faecium*. FEMS Microbes, 5, xtae018. 10.1093/femsmc/xtae018

He, Q., Hou, Q., Wang, Y., Li, J., Li, W., Kwok, L. Y., … Zhong, Z. (2018). Comparative genomic analysis of *Enterococcus faecalis*: insights into their environmental adaptations. BMC Genomics, 19(1), 527. 10.1186/s12864-018-4887-3

Hedl, M., & Rodwell, V. W. (2004). *Enterococcus faecalis* mevalonate kinase. Protein Sci., 13(3), 687–693. 10.1110/ps.03367504

Jensen, C. S., Norsigian, C. J., Fang, X., Nielsen, X. C., Christensen, J. J., Palsson, B. O., & Monk, J. M. (2020). Reconstruction and validation of a genome-scale metabolic model of *Streptococcus oralis* (iCJ415), a human commensal and opportunistic pathogen. Front. Genet., 11, 116. 10.3389/fgene.2020.00116

Jönsson, M., Saleihan, Z., Nes, I. F., & Holo, H. (2009). Construction and characterization of three lactate dehydrogenase-negative *Enterococcus faecalis* V583 mutants. Appl. Environ. Microbiol., 75(14), 4901–4903. 10.1128/aem.00344-09

Kanehisa, M., Goto, S., Hattori, M., Aoki-Kinoshita, K. F., Itoh, M., Kawashima, S., … Hirakawa, M. (2006). From genomics to chemical genomics: new developments in KEGG. Nucleic Acids Res., 34(Database issue), D354-357. 10.1093/nar/gkj102

King, Z. A., Dräger, A., Ebrahim, A., Sonnenschein, N., Lewis, N. E., & Palsson, B. O. (2015). Escher: A web application for building, sharing, and embedding data-rich visualizations of biological pathways. PLoS Comput. Biol., 11(8), e1004321. 10.1371/journal.pcbi.1004321

King, Z. A., Lu, J., Dräger, A., Miller, P., Federowicz, S., Lerman, J. A., … Lewis, N. E. (2016). BiGG Models: A platform for integrating, standardizing and sharing genome-scale models. Nucleic Acids Res., 44(D1), D515–522. 10.1093/nar/gkv1049

Lam, M. M., Seemann, T., Bulach, D. M., Gladman, S. L., Chen, H., Haring, V., … Stinear, T. P. (2012). Comparative analysis of the first complete *Enterococcus faecium* genome. J. Bacteriol., 194(9), 2334–2341. 10.1128/jb.00259-12

Levering, J., Fiedler, T., Sieg, A., van Grinsven, K. W., Hering, S., Veith, N., … Kummer, U. (2016). Genome-scale reconstruction of the *Streptococcus pyogenes* M49 metabolic network reveals growth requirements and indicates potential drug targets. J. Biotechnol., 232, 25–37. 10.1016/j.jbiotec.2016.01.035

Machado, D., Andrejev, S., Tramontano, M., & Patil, K. R. (2018). Fast automated reconstruction of genome-scale metabolic models for microbial species and communities. Nucleic Acids Res., 46(15), 7542–7553. 10.1093/nar/gky537

Magnúsdóttir, S., Heinken, A., Kutt, L., Ravcheev, D. A., Bauer, E., Noronha, A., … Thiele, I. (2017). Generation of genome-scale metabolic reconstructions for 773 members of the human gut microbiota. Nat. Biotechnol., 35(1), 81–89. 10.1038/nbt.3703

Mokhtari, A., Blancato, V. S., Repizo, G. D., Henry, C., Pikis, A., Bourand, A., … Deutscher, J. (2013). *Enterococcus faecalis* utilizes maltose by connecting two incompatible metabolic routes via a novel maltose 6’-phosphate phosphatase (MapP). Mol. Microbiol., 88(2), 234–253. 10.1111/mmi.12183

Murray, B. E. (1990). The life and times of the *Enterococcus*. Clin. Microbiol. Rev., 3(1), 46–65. 10.1128/cmr.3.1.46

O’Brien, E. J., Monk, J. M., & Palsson, B. O. (2015). Using genome-scale models to predict biological capabilities. Cell, 161(5), 971–987. 10.1016/j.cell.2015.05.019

Oberhardt, M. A., Palsson, B., & Papin, J. A. (2009). Applications of genome-scale metabolic reconstructions. Mol. Syst. Biol., 5, 320. 10.1038/msb.2009.77

Olson, R. D., Assaf, R., Brettin, T., Conrad, N., Cucinell, C., Davis, J. J., … Stevens, R. L. (2023). Introducing the Bacterial and Viral Bioinformatics Resource Center (BV-BRC): a resource combining PATRIC, IRD and ViPR. Nucleic Acids Res., 51(D1), D678–d689. 10.1093/nar/gkac1003

Paganelli, F. L., van de Kamer, T., Brouwer, E. C., Leavis, H. L., Woodford, N., Bonten, M. J., … Hendrickx, A. P. (2017). Lipoteichoic acid synthesis inhibition in combination with antibiotics abrogates growth of multidrug-resistant *Enterococcus faecium*. Int. J. Antimicrob. Agents, 49(3), 355–363. 10.1016/j.ijantimicag.2016.12.002

Pastink, M. I., Teusink, B., Hols, P., Visser, S., de Vos, W. M., & Hugenholtz, J. (2009). Genome-scale model of *Streptococcus thermophilus* LMG18311 for metabolic comparison of lactic acid bacteria. Appl. Environ. Microbiol., 75(11), 3627–3633. 10.1128/aem.00138-09

Qin, X., Galloway-Peña, J. R., Sillanpaa, J., Roh, J. H., Nallapareddy, S. R., Chowdhury, S., … Murray, B. E. (2012). Complete genome sequence of *Enterococcus faecium* strain TX16 and comparative genomic analysis of *Enterococcus faecium* genomes. BMC Microbiol., 12, 135. 10.1186/1471-2180-12-135

Ramsey, M., Hartke, A., & Huycke, M. (2014). The Physiology and Metabolism of Enterococci. In M. S. Gilmore, D. B. Clewell, Y. Ike, & N. Shankar (Eds.), Enterococci: From Commensals to Leading Causes of Drug Resistant Infection. Massachusetts Eye and Ear Infirmary.

Reed, J. L., Patel, T. R., Chen, K. H., Joyce, A. R., Applebee, M. K., Herring, C. D., … Palsson, B. O. (2006). Systems approach to refining genome annotation. Proc. Natl. Acad. Sci. USA, 103(46), 17480–17484. 10.1073/pnas.0603364103

Rice, L. B. (2008). Federal funding for the study of antimicrobial resistance in nosocomial pathogens: no ESKAPE. J. Infect. Dis., 197(8), 1079–1081. 10.1086/533452

Rychel, K., Decker, K., Sastry, A. V., Phaneuf, P. V., Poudel, S., & Palsson, B. O. (2021). iModulonDB: a knowledgebase of microbial transcriptional regulation derived from machine learning. Nucleic Acids Res., 49(D1), D112–d120. 10.1093/nar/gkaa810

Saier, M. H., Reddy, V. S., Moreno-Hagelsieb, G., Hendargo, K. J., Zhang, Y., Iddamsetty, V., … Medrano-Soto, A. (2021). The Transporter Classification Database (TCDB): 2021 update. Nucleic Acids Res., 49(D1), D461–d467. 10.1093/nar/gkaa1004

Sood, S., Malhotra, M., Das, B. K., & Kapil, A. (2008). Enterococcal infections & antimicrobial resistance. Indian J. Med. Res., 128(2), 111–121.

Staerck, C., Wasselin, V., Budin-Verneuil, A., Rincé, I., Cacaci, M., Weigel, M., … Riboulet-Bisson, E. (2021). Analysis of glycerol and dihydroxyacetone metabolism in *Enterococcus faecium*. FEMS Microbiol. Lett., 368(8). 10.1093/femsle/fnab043

Teusink, B., Wiersma, A., Molenaar, D., Francke, C., de Vos, W. M., Siezen, R. J., & Smid, E. J. (2006). Analysis of growth of *Lactobacillus plantarum* WCFS1 on a complex medium using a genome-scale metabolic model. J. Biol. Chem., 281(52), 40041–40048. 10.1074/jbc.M606263200

Thiele, I., & Palsson, B. (2010). A protocol for generating a high-quality genome-scale metabolic reconstruction. Nat. Protoc., 5(1), 93–121. 10.1038/nprot.2009.203

van der Es, D., Groenia, N. A., Laverde, D., Overkleeft, H. S., Huebner, J., van der Marel, G. A., & Codée, J. D. C. (2016). Synthesis of *E. faecium* wall teichoic acid fragments. Bioorg. Med. Chem., 24(17), 3893–3907. 10.1016/j.bmc.2016.03.019

Veith, N., Solheim, M., van Grinsven, K. W., Olivier, B. G., Levering, J., Grosseholz, R., … Kummer, U. (2015). Using a genome-scale metabolic model of *Enterococcus faecalis* V583 to assess amino acid uptake and its impact on central metabolism. Appl. Environ. Microbiol., 81(5), 1622–1633. 10.1128/aem.03279-14

Zhang, X., Rogers, M., Bierschenk, D., Bonten, M. J., Willems, R. J., & van Schaik, W. (2013). A LacI-family regulator activates maltodextrin metabolism of *Enterococcus faecium*. PLoS One, 8(8), e72285. 10.1371/journal.pone.0072285

Zhang, X., Vrijenhoek, J. E., Bonten, M. J., Willems, R. J., & van Schaik, W. (2011). A genetic element present on megaplasmids allows *Enterococcus faecium* to use raffinose as carbon source. Environ. Microbiol., 13(2), 518–528. 10.1111/j.1462-2920.2010.02355.x

Zhou, X., Willems, R. J. L., Friedrich, A. W., Rossen, J. W. A., & Bathoorn, E. (2020). *Enterococcus faecium*: from microbiological insights to practical recommendations for infection control and diagnostics. Antimicrob. Resist. Infect. Control, 9(1), 130. 10.1186/s13756-020-00770-1

Zhu, L., Zou, Q., Cao, X., & Cronan, J. E. (2019). *Enterococcus faecalis* encodes an atypical auxiliary acyl carrier protein required for efficient regulation of fatty acid synthesis by exogenous fatty acids. mBio, 10(3). 10.1128/mBio.00577-19

Zou, Q., Dong, H., & Cronan, J. E. (2024). The enteric bacterium *Enterococcus faecalis* elongates and incorporates exogenous short and medium chain fatty acids into membrane lipids. Mol. Microbiol., 122(5), 757–771. 10.1111/mmi.15322

